# Do long-tailed macaques engage in intuitive statistics?

**DOI:** 10.1101/247635

**Authors:** Sarah Placì, Johanna Eckert, Hannes Rakoczy, Julia Fischer

## Abstract

Human infants, apes and capuchins have been found to engage in intuitive statistics, generating predictions from populations to samples based on proportional information. This suggests that statistical reasoning might depend on some core knowledge that is shared with other species. Here, we investigated whether such intuitive statistical reasoning is also present in a species of Old World monkeys, to aid in the reconstruction of the evolution of this capacity. In a series of 7 test conditions, 11 long-tailed macaques were offered different pairs of populations containing varying proportions of preferred vs. neutral food items. One population always contained a higher proportion of preferred items than the other. An experimenter simultaneously drew one item out of each population, hid them in her fists and presented them to the monkey to choose. Results revealed that at least one individual seemed to make systematic population-to-sample inferences and consistently chose the sample from the population with the more favorable distribution of preferred vs. neutral food items. While it is not clear whether she used relative or absolute quantities of food, she seemed to understand the difference between a correct choice and a favorable draw and thus some basic principles of intuitive statistics.

## Introduction

The physical and social world can be described by statistical regularities: events co-occur with others repeatedly over time, resources are non-randomly distributed in space. For instance, it might rain more in some months than in others, certain fruits will be more abundant in a specific habitat, and someone’s relative will repeatedly be late at a meeting while another one will always be there in case of need. In other words, frequencies of past occurrences can be informative about existing relationships between events, as well as about the likelihood of their future occurrence. Using statistical regularities to reduce uncertainty and acquire knowledge about the state of the world, what we call statistical reasoning, is key to human learning, pervading disciplines from psychology to economics, biology, physics, law and medicine ^1–4^.

Using appropriate operations, statistical reasoning allows one to infer relationships between samples of observations and populations from which they stem. General knowledge can thus be inductively inferred from limited data^5^, and this general knowledge can in return be used to form expectations about new samples. Note that these inferences will not yield to exact predictions because of the probabilistic nature of the relation of populations and (randomly drawn) samples.

The nature and development of human intuitive statistics has long been the topic of much debate between various researchers. Some have advocated that humans become proficient in it only during later stages of childhood^6,7^ and are easily prone to make errors even as adults^8,9^, while others have argued that this ability emerges early on during childhood and plays an important role in structuring learning^10–12^. As much as an explicit understanding of probabilities and a proficient use of statistical information might be put into question, at an implicit level, there is now ample evidence that at least some aspects of statistical reasoning appear to be already present in very young children: Preverbal infants (sometime as young as 6-month-old) infer relationships between populations and samples^13^, draw inferences about physical properties of objects using statistical regularities^14^, use proportions of objects to form expectations about new samples^15^, as well as temporal and positional information of randomly moving objects to form expectations about which object was more likely to exit an urn^16,17^. In addition, they have been found to be sensitive to sampling processes and vary their expectations depending on whether the sampling appears to be random or not^18^. Whether children really engage in intuitive statistics or rather rely on simpler heuristics instead is not always clear from the data and subject to considerable debate ^7,15,17,19,20^. Nonetheless, these results, as well as results of a study of two indigenous Mayan groups^21^ show that even without formal education and language, humans appear to be intuitive statisticians.

Recently, similar reasoning abilities were highlighted in nonhuman primates: four species of apes^22^, and one species of capuchins^23^ were able to use populations of food items to form expectations about sampling events, based on a paradigm originally developed for children^15^. Another study showed that chimpanzees used proportional information to infer which of two trays containing different food/cup ratios was more likely to yield a cup containing food^24^. These studies are interesting in two aspects. Firstly, they suggest that some nonhuman primates possess some intuitive statistical abilities. Secondly, the inferences they made seemed to be based on proportions and not absolute quantities, which further points towards an understanding of some basic principles of probabilities. These findings corroborate the idea that intuitive statistics might be part of an evolutionary more ancient core knowledge that humans share with related species^25^.

The rationale of the present study was to investigate whether long-tailed macaques (*Macaca fascicularis*), a species of Old World monkeys, are able to make inferences from populations to samples, based on the same paradigm used with children^15^ and apes^22^. Adding information about a species of Old World monkeys to the emerging picture of intuitive statistical capacities in humans, great apes and capuchin monkeys, will help to further our knowledge of the distribution and potentially the evolutionary origin of these cognitive abilities within the primate order.

In a series of seven test conditions, we presented long-tailed macaques with two transparent buckets containing populations with varying proportions of preferred vs. neutral food items. Subjects watched an experimenter randomly (in appearance only) drawing a 1-item-sample out of each population and were given the choice between the two hidden samples. To receive a preferred food item as reward, therefore, subjects had to distinguish between the two populations in terms of their ratio of preferred to neutral food items, and use this relative frequency information to form expectations about the likely outcome of a sampling event. To differentiate whether monkeys really engaged in intuitive statistical inferences, or rather relied on some simpler heuristics, several control experiments were administered. Experiments 2a-c disentangled absolute and relative frequencies of food items: while one population always contained a more favorable ratio of preferred to neutral food items, the absolute quantity of preferred items was misleading (Experiment 2a) or inconclusive (Experiment 2b). Similarly, in one experiment the number of neutral food items was inconclusive (Experiment 3). Lastly, one control experiment ruled out the use of olfactory cues (Experiment 4). The underlying logic is the following: in case monkeys would engage in probabilistic reasoning, they would have a preference for the samples stemming from the populations with the higher proportion of grapes. In case they were relying on absolute number heuristics, they would have a preference for the hand drawing out of the populations with the higher quantity of grapes and/or the smaller quantity of monkey chow. If they were not expecting any quantitative information to predict sampling events, they should have no consistent preference for either population.

## Methods

### Ethical Statement

All testing was non-invasive, and subjects participated voluntarily. They were not food deprived for testing, and water was always available ad libitum. The monkeys were fed regular monkey chow, fruits and vegetables twice a day. Their enclosure was equipped with wooden platforms, fire hoses, and several enrichment objects, which were changed on a regular basis. All experiments were performed under the control of experienced veterinarians to ensure that the studies were in accordance with the NRC Guide for the Care and Use of Laboratory Animals and the European Directive 2010/63/EU on the protection of animals used for scientific purposes. In accordance with the German Animal Welfare Act, the study was approved by the Animal Welfare Officer of the German Primate Center: according to the German Law, the experiments are not invasive and do not require permission by higher authorities (LAVES Document 33.19-42502).

### Subjects

Seventeen long-tailed macaques (female N= 4) – aged 1 to 11 years (see Table 1) – participated in this study (6 of them did not reach different criteria, see supplementary materials for details). The monkeys lived in a large social group of 35 individuals. They were housed at the German Primate Center in Göttingen, Germany, and had access to indoor (49 m²) and outdoor areas (173 m²), which were equipped with branches, trunks, ropes and other enriching objects. All individuals were already experienced in participating in cognitive experiments and some of them previously took part in experiments requiring them to indicate a choice between two objects via pointing or reaching towards it. Tests were conducted once or twice a day between March and July 2016.

**Table 1.** List of subjects and conditions in which they participated.

| **Name** | **Sex** | **Date of birth** | **Participation** |
| --- | --- | --- | --- |
| Ilana | f | 07.05.2005 | Choice training |
| Paul | m | 05.06.2007 | Choice training, familiarization, Exp. 1a, 1b, 2a, 2b, 3, 4 |
| Sally | f | 17.06.2007 | Choice training, familiarization, Exp. 1a, 1b, 2a, 2b, 3 |
| Maja | f | 17.10.2007 | Choice training, familiarization, Exp. 1a, 1b, 2a, 2b, 2c, 3, 4 |
| Sophie | F | 03.04.2009 | Choice training, familiarization, Exp. 1a, 1b, 2a, 2b, 3, 4 |
| Lenny | m | 10.04.2009 | Choice training, familiarization, Exp. 1a, 1b, 2a, 2b, 3, 4 |
| Isaak | m | 10.04.2011 | Choice training, familiarization |
| Mila | f | 07.04.2012 | Choice training, familiarization, Exp. 1a, 1b, 2a, 2b, 3, 4 |
| Ilia | m | 29.12.2012 | Choice training, familiarization, Exp. 1a, 1b, 2a, 2b, 3, 4 |
| Linus | m | 16.01.2013 | Choice training, familiarization, Exp. 1a, 1b, 2a, 2b, 3, 4 |
| Max | m | 01.02.2013 | Choice training, familiarization, Exp. 1a, 1b, 2a, 2b, 3, 4 |
| Snickers | m | 12.01.2014 | Choice training, familiarization |
| Mars | m | 17.01.2014 | Choice training, familiarization, Exp. 1a, 1b, 2a, 2b, 3, 4 |
| Lord | m | 04.02.2014 | Choice training, familiarization, Exp. 1a, 1b, 2a, 2b, 3, 4 |
| Sissi | f | 07.02.2014 | Choice training, familiarization |
| Milka | f | 29.12.2014 | Choice training, familiarization |
| Sambia | f | 18.02.2015 | Choice training, familiarization |

### Experimental Setup

The testing cage (2.60 m × 2.25 m × 1.25 m; height × width × depth) was adjacent to the indoor enclosure, and could be subdivided into six experimental compartments. Subjects were tested individually in one compartment (1.05 m × 1.10 m; height × length) to which an attachable cage (73 cm × 53 cm × 35 cm; height × width × depth) was fixed, allowing subjects to have a better access to the experiment. The cage was built in metallic mesh, except for the front part that separated the monkey and the experimenter, which consisted in a removable Plexiglas pane (27 cm × 34 cm; height × length). The pane had two small holes (□ 3.5 cm; distance between holes 27 cm; see Figure 1) through which subjects could insert their arm to indicate a choice. The experimenter stood behind a wheeled table (85 cm × 80 cm × 50 cm; height × width × depth) that was set in front of the cage, on which the stimuli were presented.

**Fig. 1.**
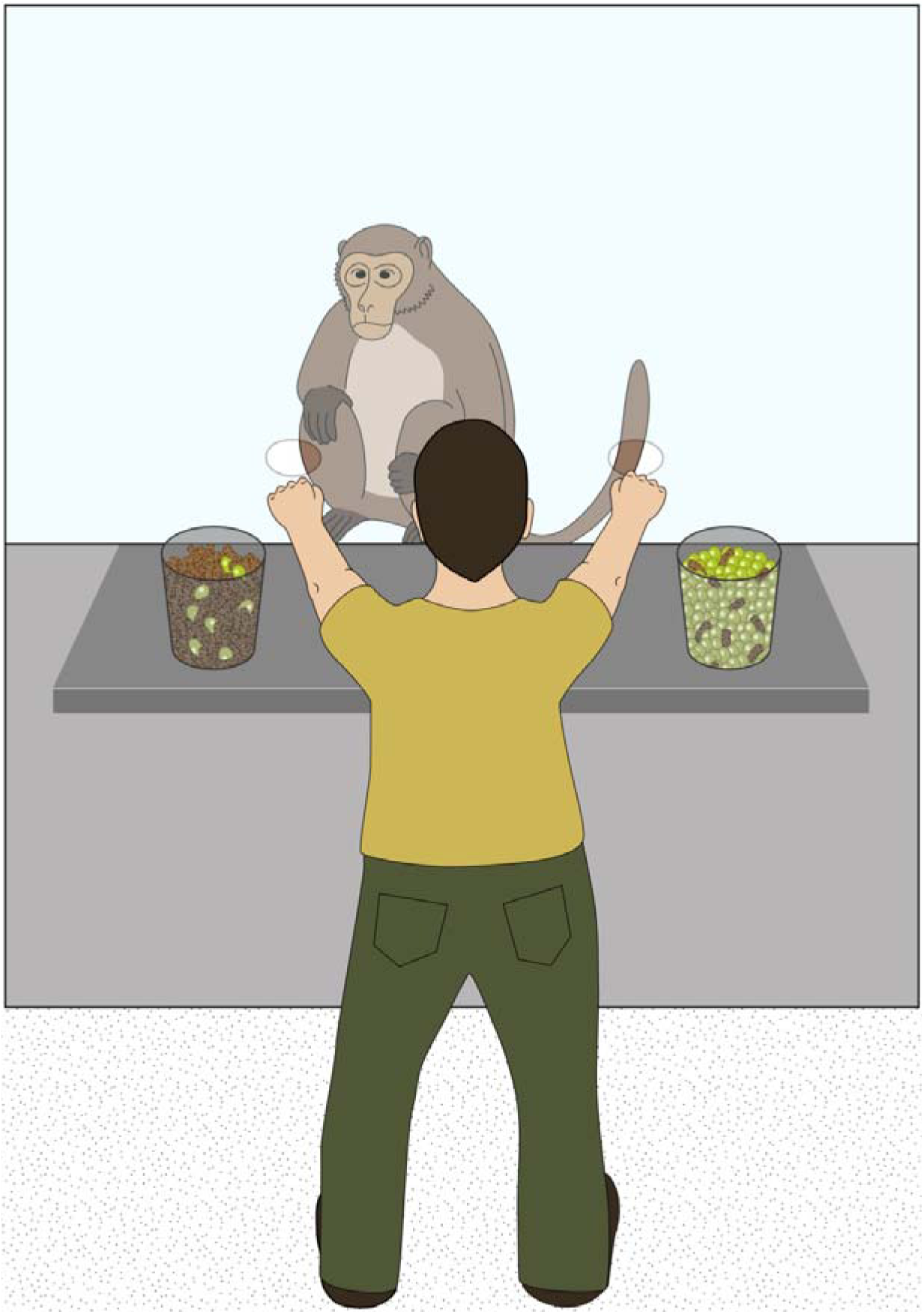
Experimental setup. The monkeys observed the experimenter drawing two hidden samples out of two populations of food items. Subsequently, the subject was given the choice between the two hidden samples.

### Study design and procedure

The study comprised four experiments (seven test conditions) preceded by a short choice training and a familiarization phase (see supplementary material for a description). Each test condition consisted of 12 test trials, evenly divided into two sessions. Test sessions always started with a preference test to make sure that monkeys´ preference for one of the two food types was consistent. Twice in a row, subjects were given the choice between one grape and one piece of monkey chow. Hence, each session consisted of two preference trials and six test trials. In each test trial, subjects were confronted with two populations consisting of a pre-determined mix of grapes (preferred food type) and monkey chow (neutral food type) contained in transparent buckets. Items of the two food types had the same approximate size but differed in color so as to render them easily distinguishable one from the other. The experimenter presented both populations on a table, shook them one after the other (always starting with the right one) and tilted them slightly forward to give the subject a good overview. She then closed her eyes, reached into the buckets and simultaneously drew one item (always the majority type, except in Experiment 3) out of each bucket in a way that kept the item hidden from the subject. While keeping the food concealed in her fists, the experimenter subsequently moved both hands towards the holes and allowed the monkey to make a choice (see Fig. 1). Once the subject had touched one of the hands, it received its content as a reward. Before the next trial started, the experimenter refilled the buckets out of sight of the subject and placed them back on the table. The position of both populations was counterbalanced across sessions and subjects (but see Exp. 1a). To make sure that subjects chose between the samples and not between the two buckets standing on the table, the experimenter crossed her arms in half of the trials before allowing the monkey to indicate a choice. Trials with and without crossing were alternated. Subjects were thus required to conclude from the information provided by the populations, which fist was more likely to contain a preferred food item as a sample.

### Coding procedure

Every session was video recorded. The experimenter coded monkeys’ choices live. Whenever monkeys chose the hand containing their preferred food item, we considered it as a success and as a failure when they chose the alternative option. A second blind observer coded 25% of the sessions, using the video recordings. Agreement between the experimenter and the second coder was perfect for all experiments (100% of agreement).

### Data analysis

To test whether monkeys’ performance as a group was different than what would be expected by chance, we computed one sample two-tailed t-tests. To analyse individual performances, we used binomial tests to calculate the probability to observe the number of successes or any higher number out of 12 trials, conditional on an underlying probability of success in one trial equal to 0.5. To adjust for an inflation of the family-wise significance level, we report p-values adjusted for multiple testing^26^. In addition, we modelled monkey´s performance conditional on whether arms were crossed or not, using a generalized linear mixed model (GLMM), with binomially distributed response and Logit link-function. The fixed effects were the two different states of the hands (crossed or not crossed). To check whether monkey learnt to associate a proportion to the preferred reward, we estimated a generalized additive mixed model (GAMM)^27^ for the successful completion of trials (binary response, Logit-link function) by allowing the probability to solve one task correctly to vary in a flexible manner, using penalized splines ^28^, i.e. such that it may potentially vary with trial, and the combination of condition and session. All these analyses were performed using R (R core Team 2015).

### Data availability statement

The datasets generated and analysed during the current study are available from the corresponding author on reasonable request.

### 3. Experiment 1a and b

In Experiment 1 we wanted to investigate whether long-tailed macaques use quantitative information to make inferences about sampling events. As Experiment 1a was not correctly randomized with regard to the side on which the hand with the preferred food appeared, we added Experiment 1b to our study, which corrected for this flaw. We used the same proportions of grapes and monkey chow in both experiments, but varied the absolute quantities of food items. Eleven subjects participated (see Table 1). Both buckets contained the same number of food items in each experiment (the total number of items was 80 in both buckets of Exp. 1a and 250 in Exp. 1b), with a distribution of grapes to monkey chow of 4:1 in one bucket and 1:4 in the other (see Table 2).

**Table 2.** Individual performance in each of the seven conditions. The proportions of grapes to monkey chow items for each population (Population A vs. Population B, grapes:monkey chow) are included for each condition. For each condition and each individual, we report the sum of correct choices within the 12 trials. * indicate performances that were statistically above chance.

| **Individual** | **Exp. 1a** | **Exp. 1b** | **Exp. 2a** | **Exp. 2b** | **Exp. 2c** | **Exp. 3** | **Exp. 4** |
| --- | --- | --- | --- | --- | --- | --- | --- |
|  | 64:16 | 200:50 | 12:3 vs. | 48:12 | 48:12 | 128:160 | 64:16 |
|  | vs. | vs. | 100:400 | vs. | vs. | vs. | vs. |
|  | 16:64 | 50:200 |  | 12:48 | 12:48 | 8:160 | 16:64 |
| Paul | 7 | 6 | 6 | 6 | / | 7 | 7 |
| Sally | 9 | 9 | 5 | 10 | / | 8 | / |
| Maja | 9 | 11* | 5 | 12* | 10 | 12* | 5 |
| Sophie | 8 | 7 | 7 | 5 | / | 7 | 7 |
| Lenny | 6 | 6 | 6 | 5 | / | 6 | 6 |
| Mila | 11* | 7 | 8 | 6 | / | 8 | 7 |
| Ilia | 6 | 6 | 5 | 7 | / | 8 | 4 |
| Linus | 6 | 7 | 6 | 6 | / | 6 | 5 |
| Max | 9 | 6 | 9 | 6 | / | 6 | 6 |
| Mars | 8 | 7 | 8 | 7 | / | 4 | 6 |
| Lord | 5 | 7 | 8 | 8 | / | 8 | 6 |

## Results and discussion

Subjects selected the sample drawn out of the favourable population in 63.6% and 59.8% of trials in Exp. 1a and Exp. 1b respectively (see Fig. 2), significantly more than expected by chance (Exp. 1a: t(10) = 3.0084, *p* = 0.01315, d = 3.01; Exp. 1b: t(10) = 2.5495, *p* = 0.02889, d = 2.55). At the individual level, only Mila and Maja were significantly above chance in Exp. 1a and 1b respectively, whereas the rest of group was not (Mila: *p* = 0.0190, Maja: *p* = 0.0159).

**Fig. 2.**
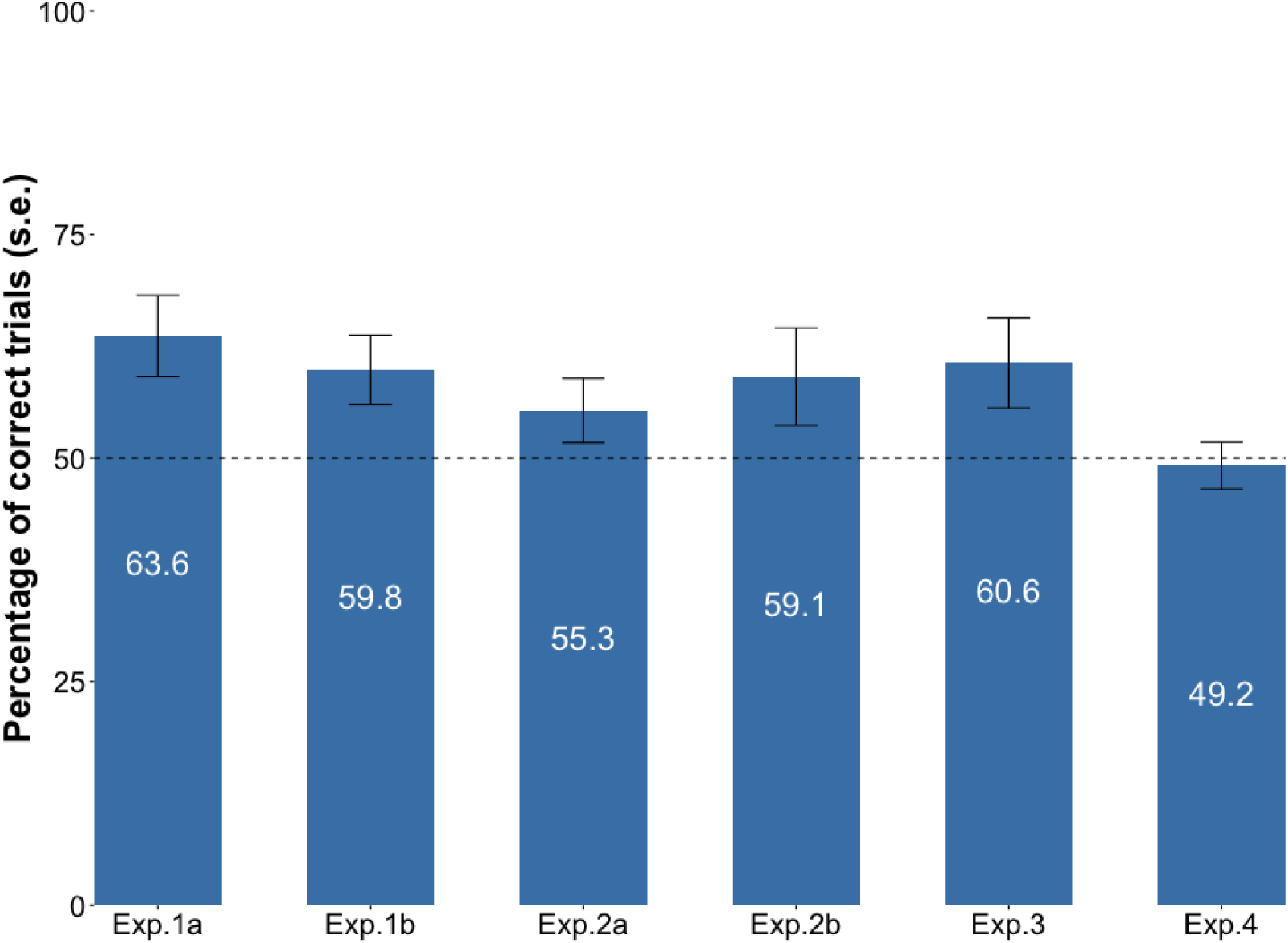
Mean percentage of trials (±1 SE) in which subjects selected the handcontaining the preferred item.

Results of Experiment 1a and b suggest that monkeys as a group were able to make inferences from populations to new samples apparently randomly drawn from the populations. At the individual level, one individual, Maja, showed the highest consistency into choosing the correct samples, followed by Sally (see Table 2). However, from this design alone it is impossible to tell whether this pattern of performance really reflects intuitive statistical reasoning or some simpler process. In particular, since the absolute and the relative frequencies of preferred vs. non-preferred food items were confounded, it is not possible to distinguish between intuitive statistics based on relative frequencies (choosing the sample with the higher probability of including a preferred food items) from merely relying on absolute frequencies (choosing the sample drawn from the population with absolutely more preferred food items). Experiments 2a-c further explores this question.

### 4. Experiment 2a, b and c

The aim of Experiment 2 was to differentiate whether subjects based their choices on the relative proportion of preferred to neutral food items, or on a simpler heuristic, namely a comparison of absolute quantities of preferred food items between both populations. Therefore, we disentangled absolute and relative frequencies of grapes in two ways: In Experiment 2a, the absolute quantity of grapes was higher in the population with the less favorable proportion of grapes to monkeys chow (see Table 2). If monkeys based their choice on the absolute quantity of preferred food items, we expected them to perform below chance level in this condition. In Experiment 2b, the absolute quantity of grapes was the same in both populations and therefore inconclusive (see Table 2). Hence, if subjects relied on absolute numbers, we expected them to choose both populations at similar rates. If, however, they used proportional information to solve the task, they should succeed in both Experiment 2a and 2b. Subjects that were successful in Experiment 2b were tested in a third condition, Experiment 2c. Experiment 2c was designed as follow up condition on Experiment 2b, to make sure that subjects recognized that both buckets contained the same absolute quantity of preferred food items. The higher quantity of neutral food items in one of the populations in Experiment 2b might have led to a visual appearance of fewer grapes in this bucket. This could allow the simple heuristic of choosing the sample from the bucket with a higher visible number of grapes. To shed light on that, we used the same quantities of food as in Experiment 2b, but this time filled the two food types in the buckets one after the other in the presence of the monkey, thereby ensuring that subjects were aware of both buckets containing the same amount of grapes. All eleven subjects participated in both Experiment 2a and 2b. Only Maja participated in Experiment 2c (see Table 1).

## Results and discussion

Subjects selected the sample drawn out of the favourable population in 55.3% and 59.1% of trials in Exp. 2a and Exp. 2b respectively, no different from chance (Exp. 2a: t(10) = 1.4725, *p* = 0.1717, d = 1.47; Exp. 2b: t(10) = 1.6705, *p* = 0.1258, d = 1.67). At the individual level, only Maja was significantly above chance in Exp. 2b (*p=*0.0017), but Sally’s performance, with 10 out of 12 correct trials, was also good (see Table 2). Maja’s performance was not above chance in Exp. 2c, but still good with 10 correct trials (see Table 2). Taken together, Experiments 2a-2c suggest that Maja and Sally did not engage in intuitive statistical inferences when widely varying numbers of grapes were used (Exp. 2a), while they seem to have used proportional reasoning when the quantity of grapes was kept constant in Exp. 2b (and 2c for Maja).

### 5. Experiment 3

The aim of this experiment was to rule out another potential alternative explanation of the patterns of results in Exp. 1. Successful performance in this experiment could have been due to truly intuitive statistics, or due to a much simpler strategy of avoiding (samples from) the population with the higher absolute number of non-preferred food items. In Experiment 3, therefore, both populations contained the same absolute number of monkey chow pieces but different amounts of grapes (see Table 2). Hence, we expected monkeys to perform at chance level in this experiment in case they relied on absolute numbers of monkey chow pellets to make their decisions. If, however, they took into account the proportion or the absolute quantity of grapes, they should succeed in this task. All eleven subjects participated (see Table 1). The procedure was the same as in Experiment 1, with the following exception: To maintain the appearance of random sampling, choosing the “correct” population did not always result in a grape as sample. Instead, sampling was proportional, i.e. the sample of Population A was a grape in five over 12 trials, with the same rewarding pattern maintained between individuals. The sample of Population B was a grape every 24 trials, meaning that half of the monkeys were never rewarded with a grape stemming from Population B. Additionally, this experiment allowed us to investigate whether monkeys would distinguish between a correct choice and a favorable outcome. Using proportions to form expectations about sampling events is one thing, understanding that because of random processes, these expectations might not match with actual outcomes is another thing ^20^. If monkeys were aware of this distinction, we predicted that their choices would be consistent throughout the 12 trials of the experiment, despite receiving neutral items as rewards for correct choices.

## Results and discussion

Subjects selected the sample drawn out of the favourable population in 60.6%, no different from chance (t(10) = 2.1058, *p* = 0.06147, d = 2.11). At the individual level, only Maja was significantly above chance (*p* = 0.0017). These results suggest that at least Maja was not relying on the absolute number of neutral items to make her inferences. Her choice was consistent across the 12 trials of the experiment, suggesting that it was not affected by the rewarding pattern.

### 6. Experiment 4

In the previous experiments, it cannot be excluded that subjects solved the task by the means of unintended cues. Experiment 4 t
herefore served as a control condition to rule out that subjects based their choice on olfactory cues or unintended cueing by the experimenter (“clever Hans”- phenomenon). Ten subjects participated (see Table 1). One female (Sally) did not enter the testing enclosure during our data collection period and was therefore not tested in this experiment. We used the same populations as in Experiment 1a (64:16 vs. 16:64) and the same procedure as in Experiment 1b, with the following exception: Both buckets were concealed by two opaque occluders, preventing subjects from seeing their content.

## Results and discussion

Results of Experiment 4 suggest that none of our subjects based their decisions on unintended or olfactory cues (t(9) = -0.318, *p* = 0.7577, d = 0.32; no individual performance was above chance).

Possible confounding effects (Exp.1 – 4)

Our results indicated no main effect of arms’ positioning, whether they were straight or crossed (Estimate + SE = 0.0574 + 0.152, z = 0.379, p-value = 0.705) Furthermore, our results showed no learning trend within and between sessions (Fig. 3).

**Fig. 3.**
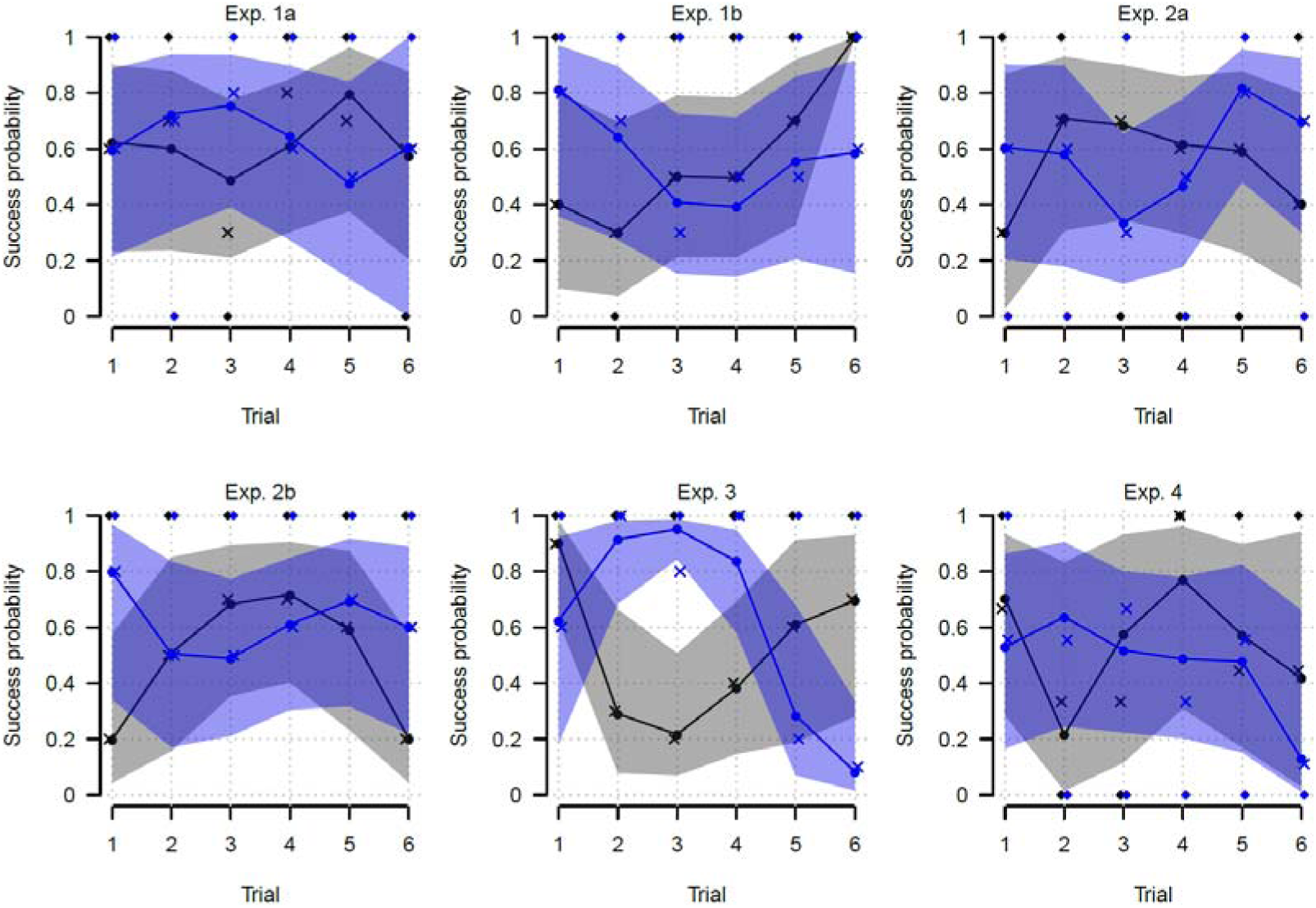
Monkeys’ performances across trials and sessions. Crosses show empirical probabilities of successfully solved trials (i.e., the number of successes divided by the number of individuals performing the trial), excluding Maja’ s performance, which is depicted by the diamonds. Shaded areas give the 99% confidence intervals for the probability of solving one trial correctly; solid lines show the estimated probabilities (both excluding Maja’s performance). Black refers to session one and blue to session 2.

## General discussion

The present findings suggest that (some) long-tailed macaques may be able to make inferences from populations to randomly drawn samples in much the same way as 12-month-old children ^15,29^, nonhuman great apes^22^ and capuchins^23^. Currently, it remains unclear, however, whether these inferences are truly intuitive statistical inferences based on relative frequencies or whether they reflect simpler heuristics that rely on information about absolute quantities. It remains also unclear how general such a capacity is given the patterns of findings at the group and individual levels. Although our group of monkeys performed above chance in both Exp. 1a and 1b, it seems that only one or two individuals were responsible for this observed pattern. These same individuals were still better than the rest of the group in most of the other experiments. In the following, we will first discuss why most individuals were at chance in our study and then evaluate Maja´s (and Sally´s) performance in more detail.

The low success rate of most of our monkeys could either reflect true competence deficits (such that intuitive statistics is simply not part of their cognitive repertoire), or they could simply point to performance deficits (though capable to do so in principle, monkeys did not recruit their intuitive statistics in the present context for some extraneous reasons).

What might such extraneous reasons and performance factors be? Possibly, individuals might have missed important steps of the procedure because they were distracted. Our results indicate that the crossing of hands was not one of those critical steps, since subjects performed on comparable levels during trials with and without crossing.

It is also possible that subjects did not pay sufficient attention to the actual quantitative information and simply attended to the presence of any grapes. Preferred food is a highly salient stimulus with potential to interfere in rational decision making ^30–32^. In a previous study, members of the same population of long-tailed macaques performed less well in a quantity discrimination task when food items served both as stimuli to be discriminated and rewards, while they performed significantly better when either inedible objects were used as stimuli or when other food items were used as rewards. Thus, the presence of grapes, which constituted both the stimuli and the (desired) reward, and which were present in both buckets, might have interfered with the comparison of proportions. To rule out this possibility, equivalent experiments with non-food items would be required. We therefore attempted to train the monkeys with black and white pebbles, representing preferred and neutral food items, but failed to bring them to criterion (data not shown).

Finally, monkeys’ competence in intuitive statistics may have been masked in the present experiments due to another performance deficit: They may simply not have bought into the premises of the task. In sampling tasks like the ones used here, predictions from the populations to the samples are justified only under the assumption that the drawing is blind. But perhaps monkeys simply did not make this assumption, either because they considered the human experimenter omniscient and almighty (not too unreasonably from their everyday experience), or because they assumed that she could haptically distinguish monkey chow from grapes during the drawing process. To address this concern, future studies could implement experiments in which the random sampling is not done by an agent but by some machine, or in which the subject herself can choose to sample. This latter possibility could also increase subjects ‘attention and make them more aware of the uncertainty linked to the outcomes and of the information needed to deal with it.

Alternatively, the present findings may reflect true competence limitations and show that macaques are incapable of using intuitive statistics (although they might still be able to engage in statistical reasoning using other formats of information, such as frequencies of sequential events for example, an ability highlighted in chimpanzees ^24^). While the present findings cannot rule out their lack of competence, one piece of evidence seems to speak against it: Maja (and Sally to a lesser extent) performed well in most conditions. This suggests that at least some of the capacities involved in statistical reasoning may be found within this species. These monkeys seemed to understand that there was a link between the composition of the populations and the drawn samples, as their decisions changed according to the food distributions. However, their performance in the critical experiments (Exp.2a and b) does not give a clear picture whether they used absolute or relative numbers of grapes. In Exp.2a, they showed no preference for either sample, whereas in Exp.2b and 2c (only for Maja) they seemed to prefer the sample from the population with the higher proportion of grapes. One explanation for this chance behavior in Exp.2a might be that the high quantity of food in Population B was too distractive for them to concentrate on proportions. This would not rule out their ability to use proportions as predictors of sampling events, but it would still show a difference with apes, as they performed above chance in a similar experiment ^22^. An alternative explanation might be that they compared information that was directly accessible to them: the absolute quantity of grapes of both populations that were visible, and not concealed by monkey chow (see Supplementary Fig. S1 of the supplementary material for pictures of the different populations). We tried to rule this alternative explanation out with Exp. 2c, in which the monkey chow was added only once Maja had seen that quantities of grapes were equal in both buckets. Her performance in this experiment, even if not perfect, was still high and suggests that she relied on proportions. To better decide between these alternatives, future studies should present information that is entirely visible to their subjects at all times.

In the initial paradigm^15^ tested on children, as well as in the two nonhuman primate studies that followed^22,23^, the rewarding pattern being always certain (at least out of the favorable population), it was not possible to assess whether subjects distinguished between a correct choice and a favorable draw. Varying the rewards following a probabilistic pattern in Exp. 3 allowed us to test whether monkeys expected outcomes to be the result of chance and irrelevant for the next choice. This seemed to be the case with Maja, as she kept choosing the correct sample despite her receiving neutral food items as reward. Maja’s performance suggests that not only was she aware that quantitative information could predict outcomes, but also that correct choices might result in unfavorable outcomes due to random processes, an idea that seems to be developing only during later stages of childhood^7,20^, but that is essential to fully understand probabilities.

In summary we found some evidence that at least one individual was making consistent inferences from populations to samples, suggesting that some intuitive statistical ability might be found within long-tailed macaques. At the group level, our subjects ‘ performance did not match the capacities described in children, great apes, and capuchins. It remains an open question whether this observed difference was due to performance limitations such as a lack of sustained attention, of motivation, or an incapacity to use a certain information format. Despite these differences, our results suggest that the ability to make inferences from populations to samples might be shared between all these species and evolutionary ancient. Maja´s general performance, as well as her consistency of choice in Exp. 3, points towards some intuitive understanding of probabilities. Nonetheless, more research is needed to better assess the information that is being used to predict sampling events (proportions or simpler heuristics), in order to further investigate the extent to which probabilities are involved in these intuitive statistical abilities. Additionally, our results stress the importance of considering individual performances and consistency across all experiments as each experiment serves as a control of another experiment. Considering only group means may lead to erroneous conclusions about group performance.

## Acknowledgments

We thank Rowan Elizabeth Titchener for her reliability assessment, Carolin Kade and Ludwig Ehrenreich for their technical assistance, all animal caretakers at the German Primate Center who assisted us and allowed us to conduct the experiments, and Holger Sennhenn-Reulen for his statistical expertise. Funding by the Deutsche 24 Forschungsgemeinschaft (GRK2070/1) is gratefully acknowledged.

## Author contributions statements

S.P, J.F and H.R conceived the study, S.P collected and analyzed the data, S.P wrote the main manuscript, and J.E, J.F and H.R reviewed it.

Authors declare no competing financial interest

## References

1. Pauker, S. G.& Kassirer, J. P. The threshold approach to clinical decision making. N. Engl. J. Med. 302, 1109–1117 (1980).

2. Bell, D. E., Raiffa, H. & Tversky, A. Descriptive, normative, and prescriptive interactions in decision making. Decision making: descriptive, normative, and prescriptive interactions, 1, 9–32 (1988).

3. Koehler, D. J. & Harvey, N. (Eds.). Blackwell Handbook of Judgment and Decision Making (John Wiley & Sons, 2008).

4. Pfannkuch, M. & Wild, C. Towards an understanding of statistical thinking. The challenge of developing statistical literacy, reasoning, and thinking. 17–46 (2004).

5. Tenenbaum, J. B., Griffiths, T. L. & Kemp, C. Theory-based Bayesian models of inductive learning and reasoning. Trends Cogn. Sci. 10, 309–318 (2006).

6. Davies, C. M. Development of the probability concept in children. Child Dev. 36, 779–788 (1965).

7. Piaget, J. & Inhelder, B. The origin of the idea of chance in children. (Trans. Leake, L., Burrel, P. & Fishbein, H.D.). (WW Norton, 1976).

8. Tversky, A. & Kahneman, D. Availability: a heuristic for judging frequency and probability. Cognitive Psychology5, 207–232 (1973).

9. Kahneman, D. & Tversky, A. On the psychology of prediction. Psychol. Rev. 80, 237–251 (1973).

10. Gopnik, A. & Wellman, H. Reconstructing constructivism: Causal models, Bayesian learning mechanisms, and the theory theory. Psychol. Bull. 138, 1085–108 (2012).

11. Xu, F. & Kushnir, T. Infants are rational constructivist learners. Curr. Dir. Psychol. Sci. 22, 28–32 (2013).

12. Xu, F. & Tenenbaum, J. B. Word learning as Bayesian inference. Psychol. Rev. 114, 245–272 (2007).

13. Denison, S., Reed, C. & Xu, F. The emergence of probabilistic reasoning in very young infants: evidence from 4.5- and 6-month-olds. Dev. Psychol. 49, 243–249 (2012).

14. Denison, S., Trikutam, P. & Xu, F. Probability versus representativeness in infancy: can infants use naïve physics to adjust population base rates in probabilistic inference?Dev. Psychol. 50, 2009–19 (2014).

15. Denison, S. & Xu, F. The origins of probabilistic inference in human infants. Cognition130, 335–347 (2014).

16. Téglás, E., Girotto, V., Gonzalez, M. & Bonatti, L. L. Intuitions of probabilities shape expectations about the future at 12 months and beyond. Proc. Natl. Acad. Sci. U. S. A. 104, 19156–9 (2007).

17. Téglás, E. et al.Pure reasoning in 12-month-olds as probabilistic inference. Science (80-.). 332, 1054–1059 (2011).

18. Gweon, H., Tenenbaum, J. B. & Schulz, L. E. Infants consider both the sample and the sampling process in inductive generalization. Proc. Natl. Acad. Sci. 107, 9066–9071 (2010).

19. Girotto, V., Fontanari, L., Gonzalez, M., Vallortigara, G. & Blaye, A. Young children do not succeed in choice tasks that imply evaluating chances. Cognition152, 32–39 (2016).

20. Falk, R., Yudilevich-Assouline, P. & Elstein, A. Children’s concept of probability as inferred from their binary choices-revisited. Educ. Stud. Math. 81, 207–233 (2012).

21. Fontanari, L., Gonzalez, M., Vallortigara, G. & Girotto V. Probabilistic cognition in two indigenous Mayan groups. Proc. Natl. Acad. Sci. U. S. A. 111, 17075–80 (2014).

22. Rakoczy, H.et al.Apes are intuitive statisticians. Cognition131, 60–8 (2014).

23. Tecwyn, E. C.,Denison, S., Messer, E. J. E. & Buchsbaum, D. Intuitive probabilistic inference in capuchin monkeys. Anim. Cogn. 20, 243–256 (2017).

24. Hanus, D. & Call, J. When maths trumps logic: probabilistic judgements in chimpanzees. Biol. Lett. 10,20140892 (2014).

25. Spelke, E. S. & Kinzler, K. D. Core knowledge. Developmental Science10, 89–96 (2007).

26. Holm, S. A simple sequentially rejective multiple test procedure. Scand. J. Stat. 6, 65–70 (1979).

27. Wood, S. N. Fast stable restricted maximum likelihood and marginal likelihood estimation of semiparametric generalized linear models. J. R. Stat. Soc. Ser. B (Statistical Methodoly)73, 3–36 (2011).

28. Wood, S. N. Thin plate regression splines Duchon spline. J. R. Stat. Soc. B65,95–114 (2003).

29. Denison, S. & Xu, F. Twelve - to 14-month-old infants can predict single-event probability with large set sizes. Dev. Sci. 13,798–803 (2010).

30. Boysen, S. T. & Berntson, G. G. Responses to quantity: Perceptual versus cognitive mechanisms in chimpanzees (Pan troglodytes). J. Exp. Psychol. Anim. Behav. Process. 21, 82–86 (1995).

31. Schmitt, V. & Fischer, J. Representational format determines numerical competence in monkeys. Nat. Commun. 2, 257 (2011).

32. Carlson, S. M.,Davis, A. C. & Leach, J. G. Less is More. Psychol. Sci. 16, 609–616 (2005).

